# Strainify: Strain-Level Microbiome Profiling for Low-Coverage Short-Read Metagenomic Datasets

**DOI:** 10.1101/2025.10.10.681738

**Authors:** Rossie S. Luo, Bryce Kille, Ellen E. Vaughan, Justin R. Clark, Anthony W. Maresso, Michael G. Nute, Todd J. Treangen

**Author notes:** Co-senior authors (Michael G. Nute, Todd J. Treangen).

## Abstract

**Motivation:** Strain-level microbiome profiling has revealed key insights into microbial community composition and strain dynamics. However, accurate strain-level analysis remains challenging due to limited linkage information, ambiguous read mapping, and complicating factors such as genome similarity, sequencing depth, and community complexity. These challenges are especially pronounced for short-read metagenomic data when estimating the relative abundances of multiple strains, a task critical for genotype-phenotype association studies.

**Results:** To address this gap, we present Strainify, which enables accurate strain-level abundance estimation from short-read metagenomes with as little as 1% genome coverage. Specifically, Strainify combines (1) identification of informative variants via core genome alignment, (2) filtering of confounding variants via a window-based test, and (3) maximum likelihood estimation of strain abundances. A Shannon entropy-weighted version of the model further improves robustness in noisy, low-coverage settings by downweighting sites with low information content. Across simulated communities of varying complexity, Strainify consistently outperformed existing approaches. On mock community sequencing data, Strainify’s estimates aligned more closely with reference abundances. When applied to a longitudinal gut microbiome dataset, Strainify successfully recapitulated the reported temporal dynamics of *Bacteroides ovatus* strain groups, demonstrating its ability to recover biologically meaningful patterns from real-world metagenomes. Together, these results establish Strainify as a robust and versatile solution for accurate strain-level abundance estimation in short-read, low-coverage microbiome studies.

**Availability:** The Strainify code and results are available at: https://github.com/treangenlab/Strainify

## Introduction

Recent advances in microbiome sequencing have led to an increased interest in strain-level comparative genomic analysis and the exploration of genotype–phenotype relationships, particularly within highly complex microbial communities such as the human gut microbiome^1,2^. Strain diversity has emerged as a key factor in bacterial pathogenicity and drug–microbe interactions^3^. Evidence has shown that various species, such as *Escherichia coli* and *Bacteroides fragilis* have high intraspecific functional diversity and can be classified as either pathogenic or beneficial depending on fine-grained genetic differences^4–6^. For example, a commonly used probiotic *E. coli* strain Nissle 1917 was found to also display mutagenic activity via the colibactin-producing operon, which could potentially induce colorectal cancer^7^. Moreover, specific biological pathways involved in microbial drug metabolism can be significantly influenced by strain-level differences, in some cases by a single nucleotide polymorphism (SNP)^3^. In addition, microbiome studies investigating mother-to-infant transmission^8,9^and fecal microbiota transplants (FMT)^10,11^ have underscored the importance of characterizing strain-level composition to understand microbial inheritance and community dynamics. Strain-resolved analyses further reveal that ecological and clinical factors, such as propagule pressure and resource competition govern donor strain engraftment after FMT^12^. Overall, these findings highlight the critical need for accurate strain-level analysis to uncover functional and ecological variation that would otherwise be obscured at the species level^3^.

Numerous computational approaches have been developed to estimate species-level composition from metagenomic data. For example, MetaPhlAn 4^13^ uses a marker-based strategy, whereas TIPP3 combines phylogenetic placement with abundance estimation to achieve accurate and scalable species-level profiling in complex microbial communities^14^. However, strain-level profiling methods are needed to resolve fine-scale diversity *within* species. Variants that distinguish closely related strains often lie outside marker regions, and strain-level tools must consider all possible variants, since only a few discriminatory mutations may exist, especially in the context of genotype-to-phenotype analyses or competitions. While strain-specific genomic elements can reveal the presence or absence of strains, they alone are insufficient to accurately quantify relative abundances. Estimating relative abundance is a crucial step in computing differential enrichment, which in turn enables the association of taxonomic identities or genotypes with experimental conditions and ultimately phenotypic outcomes^15,16^. Quantitative strain composition analysis is complicated by several factors, including sequencing depth, number of strains, unevenness of abundance distributions, and degree of genetic divergence between strains^15^. Existing tools take various approaches to address these challenges and generally fall into two categories: (1) reference-based methods, which typically provide higher specificity, and (2) assembly-based methods, which tend to achieve higher sensitivity but require sufficient and relatively uniform genome coverage to accurately reconstruct individual strains^15,17^. However, few tools have achieved both high sensitivity and specificity. As a result, the reliable detection of low-abundance strains remains a major obstacle, and their underrepresentation or absence in assembled genomes can lead to systematic underestimation of key biological signals and misinterpretation of microbiome dynamics^18^.

Short-read metagenomic sequencing is a popular approach in microbiome studies due to its cost-effectiveness and efficiency^19^. However, it presents a set of unique challenges compared to other sequencing methods. The limited read length (150 base pairs on average) makes read mapping and assignment difficult at strain-level where genomes are highly similar with an average nucleotide identity above 99.99%^19,20^. Many existing strain-level profiling tools adopt marker- or gene-based approaches. For example, StrainPhlAn^21^ identifies and maps reads to species-specific marker genes while ConStrains^22^ infers within-species structures by analyzing SNP patterns in a set of universal genes. PanPhlAn^23^ extends these approaches by using a species’ pangenome to identify strain-specific gene presence–absence patterns. Similarly, DESMAN infers strain abundances and haplotypes from variant frequencies in core genes of metagenome-assembled genomes^24^. These approaches offer limited resolution as markers may not exist when strains share an almost identical core genome with only a small number of SNPs. In addition, low sequencing depth which is common in complex microbial communities or in microbiome samples with a majority of host DNA, can significantly increase the difficulty of parsing signal from noise in short reads. A comparison of existing strain-level profiling tools for short-read metagenomic data is summarized in Table 1.

**Table 1.**
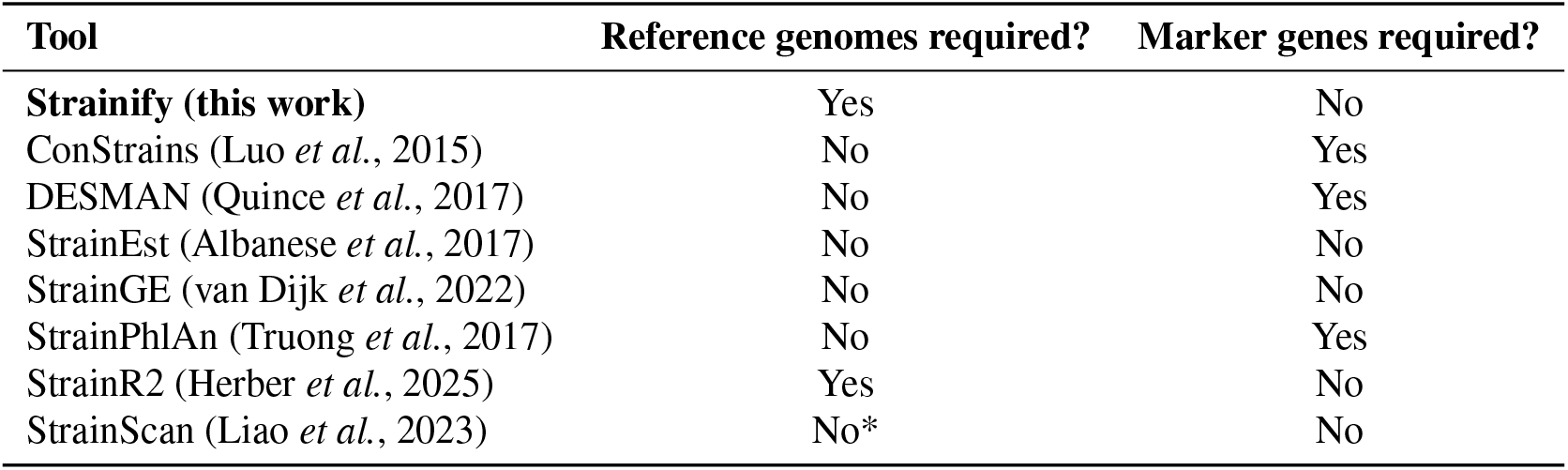
Comparison of existing strain-level profiling tools for short-read metagenomic data. *Reference genomes required?* – whether external reference genomes must be provided (or improve accuracy). *Marker genes required?* – whether detection relies on a pre-defined marker gene set. *StrainScan’s accuracy improves with custom database built with known references.

| Tool | Reference genomes required? | Marker genes required? |
| --- | --- | --- |
| **Strainify (this work)** | Yes | No |
| ConStrains (Luo *et al.*, 2015) | No | Yes |
| DESMAN (Quince *et al.*, 2017) | No | Yes |
| StrainEst (Albanese *et al.*, 2017) | No | No |
| StrainGE (van Dijk *et al.*, 2022) | No | No |
| StrainPhlAn (Truong *et al.*, 2017) | No | Yes |
| StrainR2 (Herber *et al.*, 2025) | Yes | No |
| StrainScan (Liao *et al.*, 2023) | No* | No |

To address these challenges and enable high-precision microbiome discoveries, we developed Strainify, a robust strain-level abundance analysis tool for short-read metagenomic data with both high sensitivity and specificity, even in low-coverage samples. Strainify applies a maximum likelihood model (MLE) that leverages allele frequencies *across the genome*, including sites that do not individually distinguish strains, to infer the vector of abundances that best explains the data. Informative variant sites are identified from a core genome alignment, while variants likely influenced by recombination are excluded to reduce bias. Strainify achieves accuracy typically within 1% of ground truth, even on mixtures of 30 strains at average coverages below 1.0×. Using simulations and a mock community, we demonstrate that Strainify is robust to skewed abundance distributions, low coverage, and variation in microbial species. Furthermore, Strainify accurately recapitulated strain-level dynamics in a longitudinal gut microbiome study, underscoring its power to resolve population structure with high precision and demonstrating its effectiveness in real-world applications. By enabling accurate strain-level abundance estimation, Strainify provides a powerful framework for investigating microbial community dynamics in diverse environments and for uncovering their implications for human health and disease.

## Results

To test the performance of Strainify, we conducted multiple experiments which involved both simulated and real datasets. In all experiments conducted with simulated or mock data, Strainify (the updates branch of the Strainify repository) was benchmarked against StrainScan (V1.0.14), a recently published tool designed specifically for strain-level relative abundance analysis^25^. StrainScan constructs a matrix of strain-specific k-mers, counts their occurrences in the sample and uses matrix multiplication to estimate the relative abundance vector. By contrast, Strainify builds a variant matrix based on SNPs and indels in the core genome and infers the abundance vector using a maximum likelihood framework. Although the underlying features and inference models differ, both methods rely on matrix representations to resolve strain-level composition from information pulled from sites across the genome, so StrainScan is a highly relevant point of comparison. In the mock dataset, we also included StrainR2 (V2.3.0)^26^, another recent tool that estimates strain abundances by mapping reads to reference genomes and applying normalization based on unique k-mers from each genome. However, StrainR2 was excluded from simulated benchmarking experiments due to its failure to produce results from 250 bp reads. The StrainR2 paper reported results for reads up to a maximum length of 150 bp^26^.

### Simulated dataset with 4 *E*.*coli* strains

We simulated a mock community which consisted of the same 4 *E. coli* strains used in the StrainScan manuscript: E24377A, H10407, Sakai, and UTI89. The genomes were obtained from NCBI. The strains are taken from a mock community with publicly available short-read sequencing data (SRR13355226), and that dataset is also used to evaluate the methods.

To demonstrate the effect of having different shapes in the underlying abundance distribution, four different abundance vectors (Table 2) were simulated to mimic the following scenarios: 1) log skewed, with abundances varying by multiple orders of magnitude (“log”), 2) uniform, 3) asymmetric, and 4) asymmetric (reversed). The last of these is intended to test whether the outcome on the asymmetric distribution may depend on which genome is higher or lower. Three coverages were simulated for each ratio: 10×, 20×, 50×. Higher coverage levels were also simulated, though the results were substantially similar to 50×. ART^27^ was used to simulate paired-end Illumina short reads. For each of the coverages, reads from each genome were simulated separately with the following parameters: `-ss MSv3 -p -l 250 -f IC -m 600 -s 150`, where `IC` (for individual coverage) is equal to the total coverage multiplied by the proportion of each genome in each abundance ratio. The reads from all four genomes were then mixed to create simulated metagenomes for abundance analyses.

**Table 2.** Abundance ratios in simulated *E. coli* metagenomic samples.

| Genomes | log | uniform | asym 1 | asym 2 |
| --- | --- | --- | --- | --- |
| E24377A | 0.001 | 0.25 | 0.1 | 0.4 |
| H10407 | 0.800 | 0.25 | 0.2 | 0.3 |
| Sakai | 0.049 | 0.25 | 0.3 | 0.2 |
| UTI89 | 0.150 | 0.25 | 0.4 | 0.1 |

Strainify was compared against both the default and low coverage settings of StrainScan. A custom database was constructed for StrainScan using the query genomes. The `-l` parameter was set to 2 in the low coverage setting of StrainScan which is recommended for samples with less than 1× coverage. For Strainify, variant filtering window size was set to 500 base pairs.

Figure 2 compares the absolute error between each tool and the ground truth, by taxon, by dataset. Strainify demonstrated a clear advantage over StrainScan, particularly at lower coverages. Strainify has lower absolute error on nearly every taxon and nearly every genome with the exception of E24377A on the uniform abundances. Note that for this particular distribution, the y-axes of these panels go from -1% to 2% whereas panels in all other columns have a range of at least +/- 5%. On nearly every other taxon, coverage and abundance distribution, Strainify produces more accurate estimates. Overall, Strainify achieved an average absolute error of just 1.016%, which is 3 times lower compared to default StrainScan (3.079%) and 4 times lower compared to low coverage StrainScan (4.054%).

**Figure 1.**
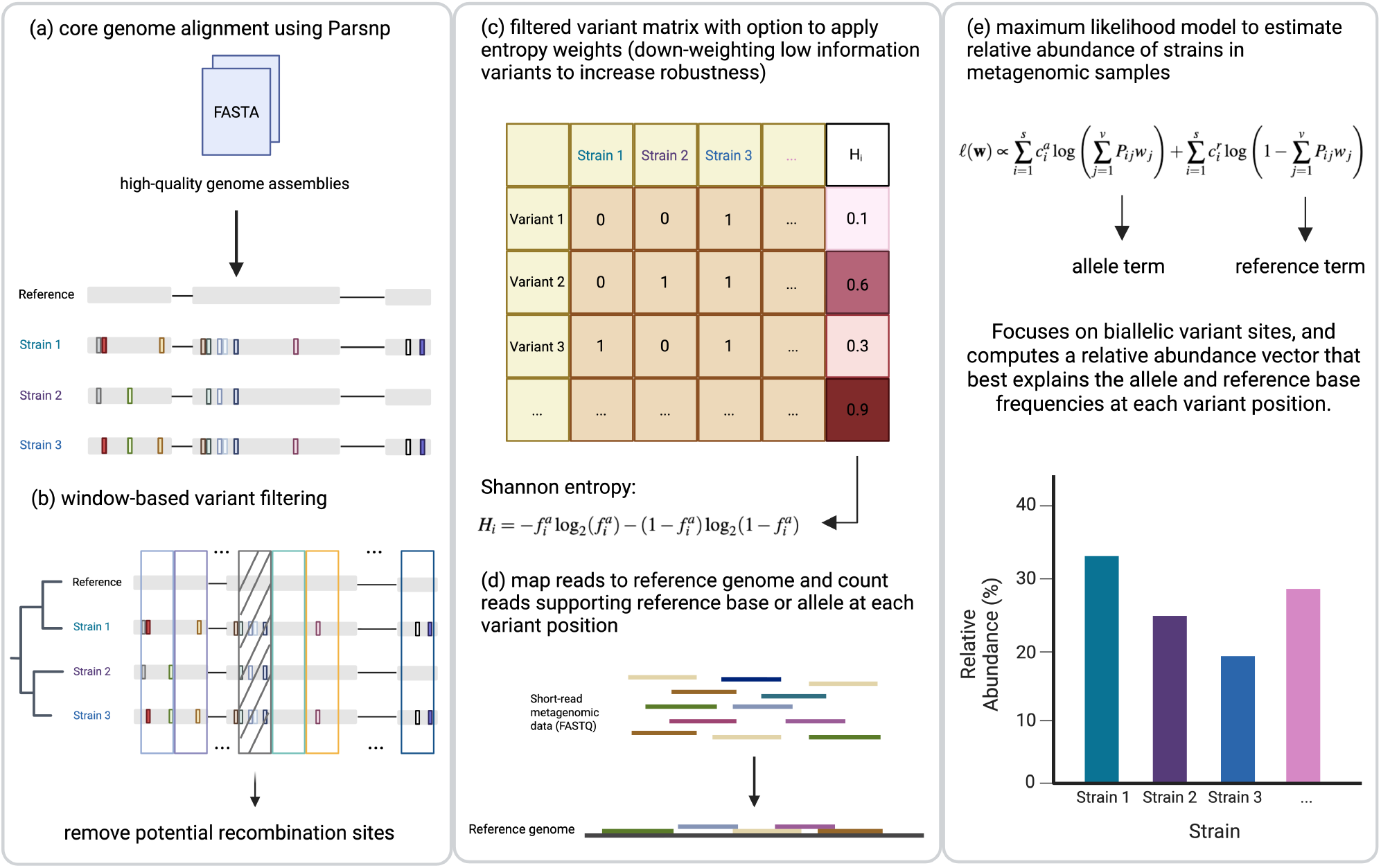
Strainify workflow. (a) Core genome alignment is performed with Parsnp on high-quality assemblies of strains expected in the metagenomic samples. (b) The core genome is partitioned into windows, and a chi-squared test is applied to remove those with significantly higher variant density, indicative of recombination. (c) A variant-by-strain matrix is generated, with optional Shannon entropy weighting to down-weight low-information variants under noisy conditions. (d) Short reads are mapped to the same reference genome used in step (a), and read counts are obtained at variant positions defined in step (c). (e) A maximum likelihood model estimates relative strain abundances. It focuses on biallelic variant sites and optimizes a vector of abundances that best explains the observed allele and reference frequencies.

**Figure 2.**
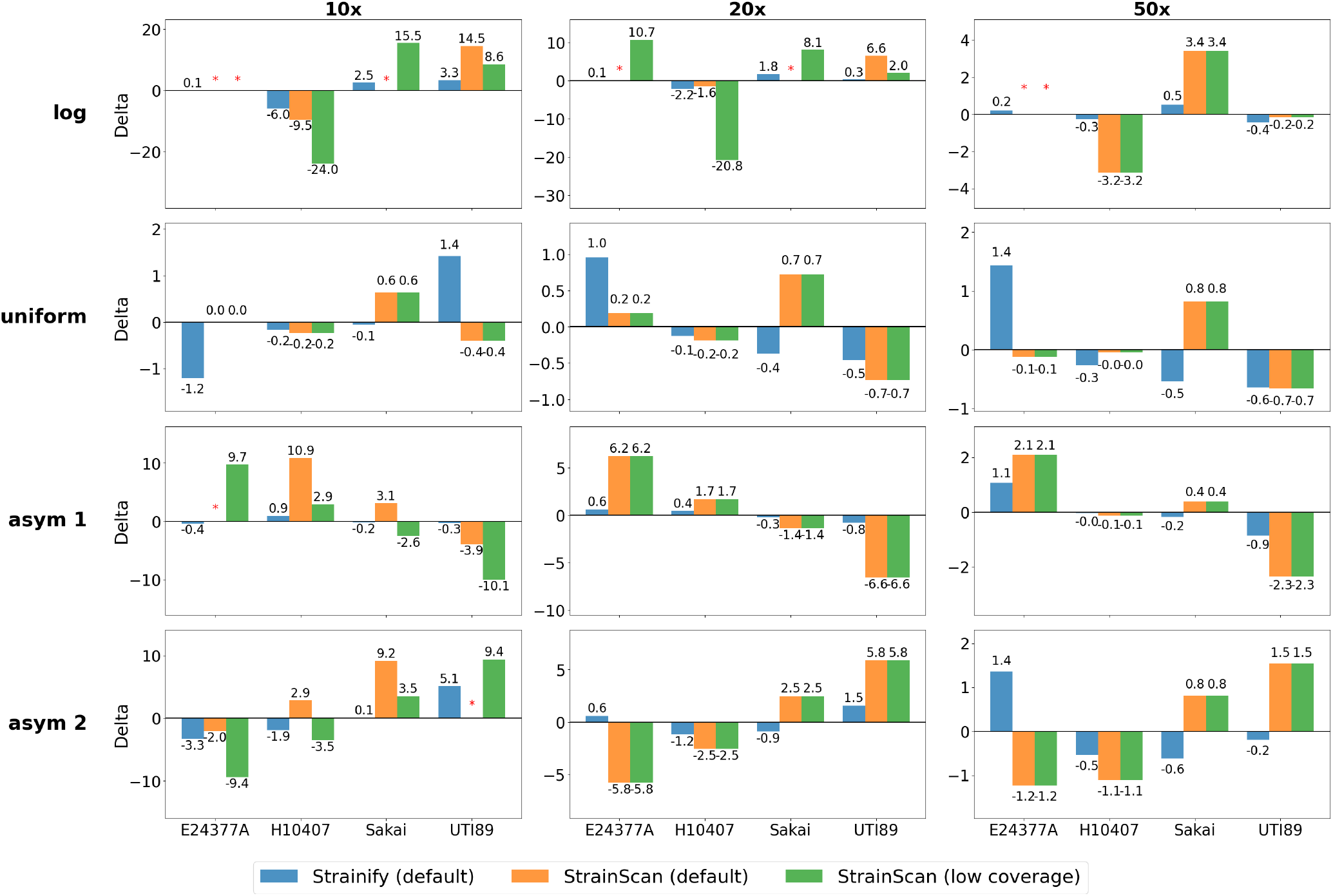
Comparison of Strainify and StrainScan on a simulated dataset of 4 *E. coli* strains. Delta = estimate (%) - ground truth (%). Larger deviations from 0 indicate greater divergence from the true abundance. Asterisk (*) indicates that the strain was undetected. The log distribution includes a dominant strain (80%) and a rare strain (0.01%). In the uniform distribution, all strains have equal abundance. Asym 1 and asym 2 represent abundance gradients in ascending and descending order, respectively.

The first column of panels is particularly noteworthy because StrainScan appears to struggle at extremely low coverage levels. In the 10× log distribution (top left panel), the low-abundance strain (E24377A) is present at just 0.01% abundance. At these ultralow abundances, StrainScan does not provide reliable estimates. Its low-coverage mode introduces substantial error by biasing the abundance estimate of the dominant strain. Across the distributions, there are cases where default StrainScan performs well but low-coverage StrainScan performs poorly, and vice versa, often within the same dataset. In the 20× log distribution data (second panel of the first row), E24377A, still present with only 0.01% abundance, is correctly identified and quantified by Strainify, while the default StrainScan does not detect it and the low-coverage StrainScan setting significantly overestimates its abundance. This highlights the particular difficulty of accurately handling strains at very low coverage, a challenge that Strainify is able to overcome.

### Simulated datasets with 30 strains

To assess the performance of Strainify in more complex metagenomic samples containing a larger number of strains, we simulated datasets with 30 strains from the same species. Five species (*C. acnes, C. difficile, E. coli, M. tuberculosis* and *S. epidermidis*) were selected for this experiment. *C. acnes, C. difficile, E. coli* and *S. epidermidis* are bacteria commonly found in skin or intestinal microbiome and can potentially cause severe infections. *M. tuberculosis* is the organism responsible for the airborne infectious disease tuberculosis (TB) and is known to have high genome similarity between strains^28^, which may present computational challenges in strain-level abundance analysis.

The large number of strains in the single experiment here is intended to test the ability to deconvolve this many strains. The challenge of low abundance and low coverage is amplified not only because, by necessity, nearly all abundance values are low, but also because, on a relative scale, small absolute errors can represent large proportional inaccuracies. For example, in a uniform distribution, a 1% absolute error corresponds to a 30% relative error compared to the ground truth abundance.

For each species, 30 genomes were randomly selected and retrieved from NCBI. These genomes were then used to generate simulated metagenomic samples with three abundance ratios. In abundance ratio 1, one dominant strain was randomly selected out of the 30 strains and accounts for 80% abundance (“dominant”). The remaining 20% abundance was then randomly distributed among the other strains. Abundance ratio 2 simulated an even distribution (“uniform”) while abundance ratio 3 simulated a distribution which was itself simulated from a Dirichlet-Uniform distribution (“dirichlet”). Five coverages were included: 10×, 20×, 50×, 100× and 200×. ART^27^ was used to create the simulated metagenomic datasets with the same settings from the experiment described above. Strainify was again benchmarked against StrainScan’s default and low coverage settings. In addition to Strainify’s default mode, its entropy weighting mode was also included in the comparison since higher noise is expected as the number of strains to de-convolute increases.

As shown in Figure 3, Strainify outperformed StrainScan across all five strains, achieving a Jensen-Shannon Divergence (JSD) consistently below 0.1 across all abundance ratios and sequencing coverages. Strainify improved performance is particularly evident at lower coverages, such as 10× and 20×, where StrainScan’s performance degraded due to increased noise. This effect is most pronounced in the especially challenging scenario of *M. tuberculosis* at 10× coverage, where the sample includes highly similar genomes and low sequencing depth. Strainify is able to maintain high accuracy while StrainScan has a sharp spike in its JSD. StrainScan’s results at low coverage are mainly due to limited detection of low-abundance strains. For example, StrainScan’s default mode only detected 5 out of 30 strains in ratio 2 of the *M. tuberculosis* dataset with 10× coverage. In contrast, Strainify successfully detected all 30 strains in this dataset.

**Figure 3.**
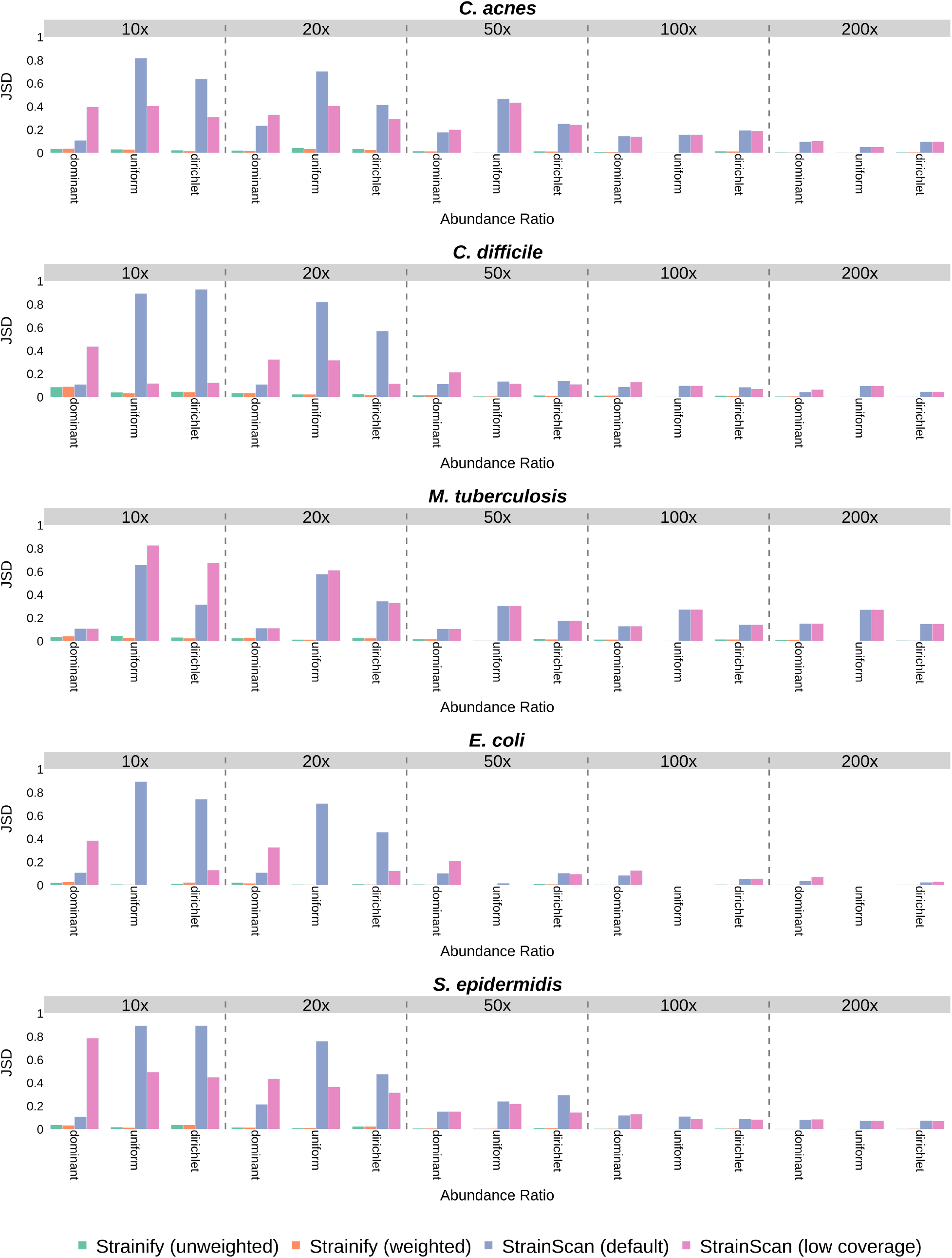
Jensen-Shannon Divergence (JSD) of Strainify and StrainScan between ground truth and estimated relative abundance of a simulated dataset consisting of 30 strains across five species (*C. acnes, C. difficile, M. tuberculosis, E. coli* and *S. epidermidis*).

In this experiment, entropy weighting of variants helped the embedded conic solver (ECOS) converge more reliably and reduced dependence on the backup solver. For instance, in one dataset (*E. coli*, 20× coverage, ratio 1), ECOS failed due to numerical instability and the splitting conic solver (SCS) was used. In this case, the entropy-weighted model improved both convergence and accuracy, yielding results closer to the ground truth.

### Real dataset from a mock community of 4 *E. coli* strains

To further validate Strainify’s performance on real datasets, we downloaded a dataset used in the StrainScan paper (SRR13355226). This dataset contains paired-end Illumina reads from a mock community with 99% human DNA and 1% *E. coli* DNA. The mock community consists of the same four *E. coli* strains used in the earlier simulated 4-strain experiment. Strainify’s estimates were compared against the published benchmarking results from the StrainScan paper (including results from StrainScan, StrainGE and StrainEst), results from StrainR2, and a reference ground truth (Figure 4). The reference ground truth is identical to the “log” abundance distribution in the simulated 4-strain experiment. In this context, the “ground truth” does not reflect the actual abundances of the strains in the experimental mixture, but rather serves as a reference point to facilitate comparison across methods.

**Figure 4.**
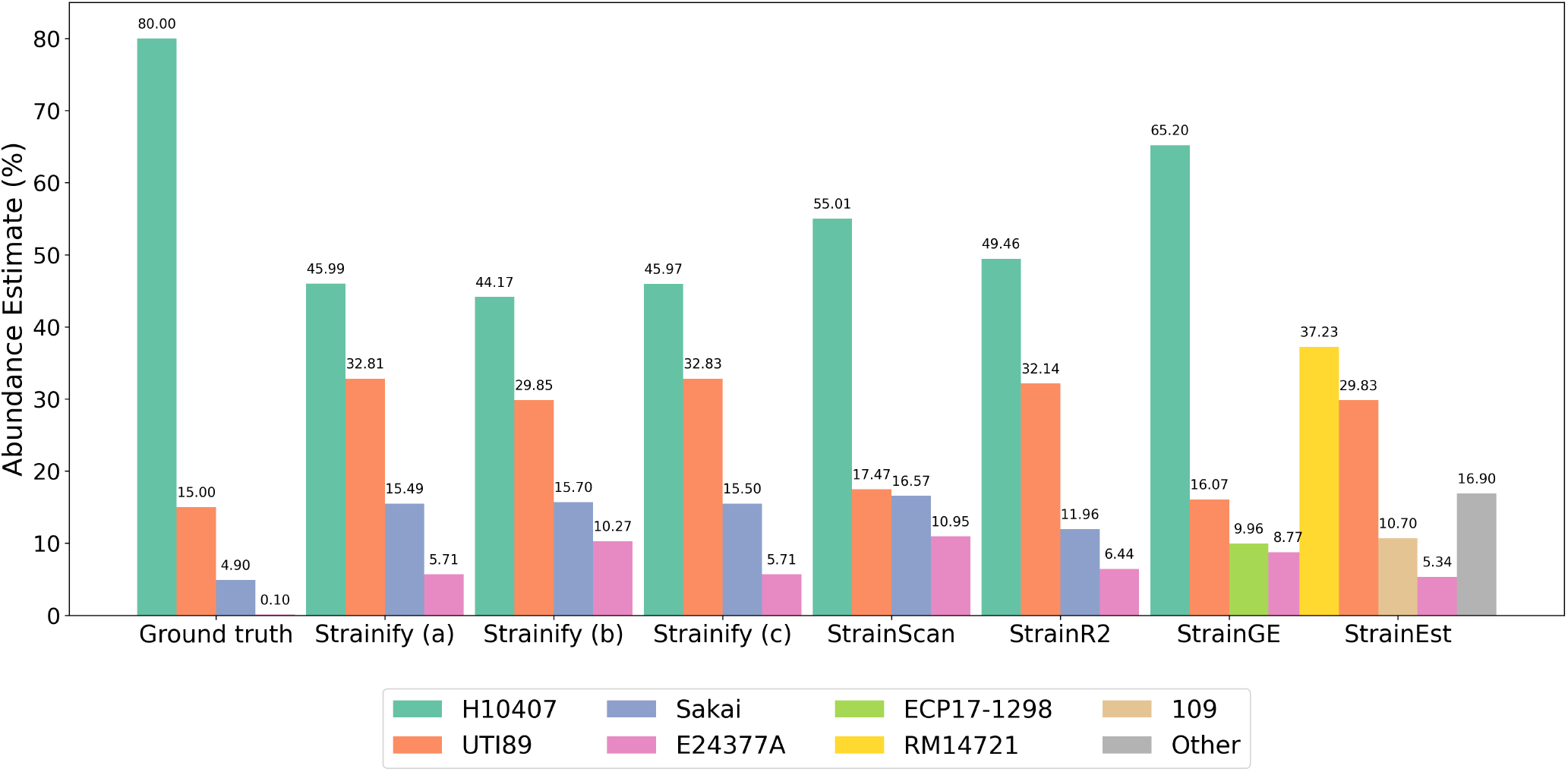
Comparison of relative abundances (%) in a mock community of four *E. coli* strains. Estimates were obtained using Strainify under three conditions: (a) plasmids included, (b) plasmids excluded entirely, and (c) plasmids included only for genome alignment and read mapping. For comparison, results from StrainScan, StrainR2, StrainGE, and StrainEst are also shown. Data for StrainScan, StrainGE, and StrainEst were obtained from the StrainScan publication. “RM14721” serves as the representative strain for “H10407” in StrainEst, while “ECP17-1298” and “109” represent “Sakai” in StrainGE and StrainEst, respectively^25^. The reference “ground truth” corresponds to the log distribution reported in Table 2, representing the intended abundance ratios of strains manually mixed to construct the mock community. Note that this ground truth does not reflect the source ratio of sequencing reads in the metagenomic sample.

Since all four strains contain plasmids, which may influence the abundance estimates when plasmid concentrations are not proportional to the abundances of the strains, we evaluated the effect of excluding plasmids both entirely and partially in Strainify’s analysis pipeline. For entire exclusion, plasmids were removed from the input FASTA files while for partial exclusion, plasmids were included only in the genome alignment and read mapping steps but not the rest of the pipeline.

The estimated abundance rankings of the four strains were consistent across Strainify, StrainScan, StrainR2, StrainGE and StrainEst. Strainify and StrainR2 produced highly similar results. Compared to StrainScan, StrainGE and StrainEst, Strainify’s estimates more closely aligned with the reference ground truth, showing a clearer separation between strains. For instance, the ground truth indicates that UTI89 is approximately three times more abundant than Sakai. Strainify estimated a ratio of roughly 2:1, while StrainScan produced nearly equal estimates for the two strains, with only a slight increase for UTI89. Similarly, StrainGE failed to clearly distinguish between ECP17-1298 (the representative strain of Sakai) and E24377A. In contrast, StrainEst assigned a substantial fraction of the abundance to strains not present in the dataset, leading to an underestimation of the most abundant strain. These results demonstrate that Strainify performs competitively with other state-of-the-art tools.

Partial exclusion of plasmids had little effect on Strainify’s estimates. However, complete exclusion altered the estimated ratio between E24377A and Sakai from approximately 1:3 to 1:1.5, further diverging from the reference ground truth ratio of 1:49. This suggests that excluding plasmids entirely may introduce bias and is likely unnecessary in this context.

### Longitudinal dataset of *Bacteroides ovatus* in human gut microbiome

Tracking strain-level abundance of specific species in the gut microbiome over extended periods provides valuable insights into key microbial community members and their potential impacts on host health, including disease associations. However, real-world metagenomic samples are challenging to analyze due to their complex microbial composition and potential contamination. To address this, we evaluated Strainify’s ability to estimate strain-level abundances in longitudinal human gut microbiome samples.

We applied Strainify on a dataset from a longitudinal gut microbiome study^29^ in which stool samples were collected from a fecal microbiota transplant donor and the relative abundances of *Bacteroides ovatus* strains were tracked for 540 days. A total of 89 fully assembled *B. ovatus* genomes from this study with completeness above 90% were downloaded from NCBI and used as one of Strainify’s inputs. Strainify was run in entropy-weighting mode as noise was expected to be high in these complex, multi-strain samples.

For direct comparison to the original study, we adopted the same phylogeny and strain groupings, and aggregated strain-level estimates into the defined groups. As shown in Figure 5, Strainify recovered highly similar temporal abundance patterns to those reported previously. For example, strain group 2 initially dominated but was gradually overtaken by strain group 3 within the first 200 days. Strain group 1 remained the least abundant throughout the study, although it increased in relative abundance between days 200 and 400. These results demonstrate that Strainify can robustly reveal biologically meaningful strain-level dynamics from complex longitudinal gut microbiome datasets, underscoring its utility for studying microbial population structure in real-world metagenomic samples.

**Figure 5.**
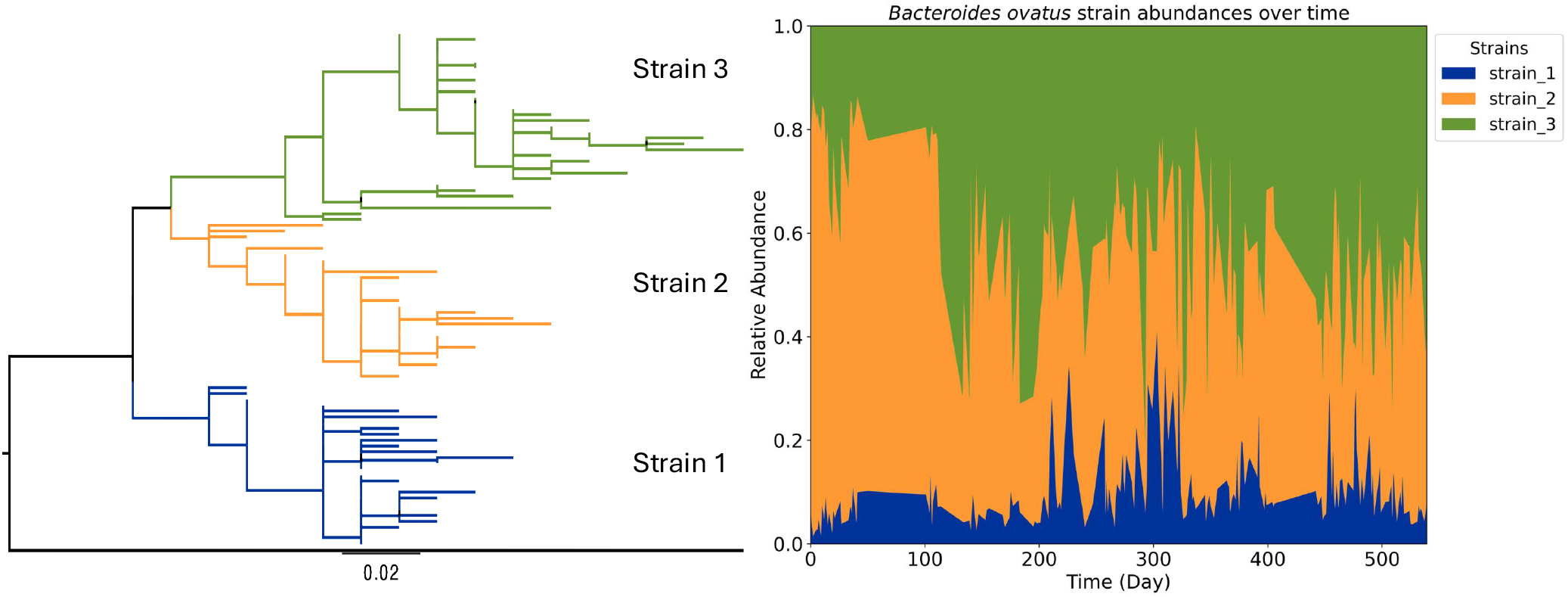
Relative abundances of *Bacteroides ovatus* strains in a longitudinal human gut microbiome study. Strains were grouped into three main strains based on the phylogeny tree from Poyet *et al*., 2019^29^. Strains with no fully assembled genomes available on NCBI or with low genome assembly completeness (lower than 90%) were excluded from the analysis.

### Runtime and memory usage

All benchmarking experiments were performed on an x86_64 Linux server equipped with 80 Intel® Xeon® Gold 6138 CPUs running at 2.00 GHz. In all experiments, Strainify was executed using 12 threads for both the *build variant matrix* and *compute abundances* steps. StrainScan was also configured to use 12 threads for the *build database* step. However, it does not allow user control of the number of threads during the *compute abundances* step. Similarly, StrainR2 was executed with its default configuration, which does not provide user control of the thread parameter. Wall clock time (s) and peak RAM (MB) for Strainify were obtained using Snakemake’s internal benchmarking feature (based on `psutil`^30^), while the same metrics for StrainScan and StrainR2 were collected using `/usr/bin/time -v`. Comparisons of runtime and memory between Strainify and StrainScan for the 4-strain simulated dataset and the 30-strain simulated dataset are presented in Supplementary Figures 1–4.

Overall, Strainify was highly efficient in the *build variant matrix* step, requiring less than 15 minutes and 2.5 GB of memory across datasets of varying sizes and microbial community complexity. In contrast, StrainScan required substantially longer runtime and higher memory usage to build its database as the number of strains and community complexity increased. In particular, in the 30-strain experiment, it required over 2 hours to build the *C. difficile* database and approximately 30 GB of memory to build the *E. coli* database (Supplementary Figure 2). For the *compute abundances* step, Strainify required up to 15 minutes and 1.5 GB of memory per sample, whereas StrainScan required up to 3.5 minutes and 2 GB of memory per sample (Supplementary Figures 1, 3 and 4). When compared alongside both StrainScan and StrainR2 in the 4-strain *E. coli* mock community experiment (Figure 6), although Strainify had higher wall clock time than StrainR2, it demonstrated the smallest memory footprint and an end-to-end runtime comparable to that of StrainScan.

**Figure 6.**
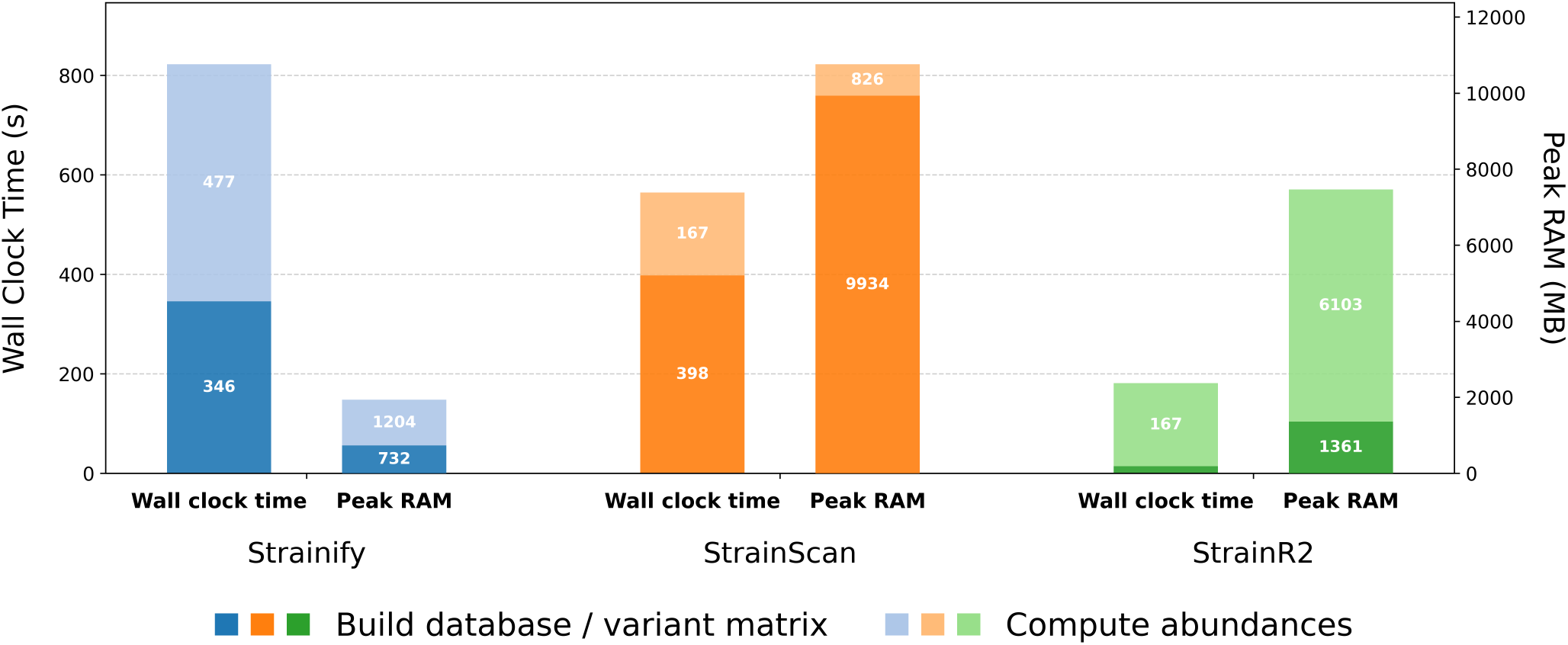
Wall clock time (s) and peak RAM (MB) of Strainify, StrainScan, and StrainR2 in the 4-strain *E. coli* mock community benchmarking dataset (SRR13355226). Default settings were used for all three tools. Both wall clock time and peak RAM are measured in two stages: (i) the *build database* step for StrainScan and StrainR2, and the corresponding *build variant matrix* step for Strainify, and (ii) the *compute abundances* step.

## Discussion

Overall, Strainify achieved high accuracy across a broad range of sequencing coverages and abundance distributions, consistently maintaining JSD values below 0.1. Strainify demonstrated improved performance compared to a leading tool for strain-level microbiome analysis in StrainScan, particularly at low coverages. While StrainScan’s performance declined substantially when individual strain coverages dropped below 1×, Strainify accurately recovered strain-level abundance profiles even at extremely low genome coverage levels, down to only 1% of the genome covered. Strainify’s entropy-weighted MLE model performed well in challenging scenarios characterized by sparse data and high noise by down-weighting low-information variants. Furthermore, Strainify exhibited strong performance in cases of genomic divergence, as evidenced by its ability to correctly identify and quantify all closely related *M. tuberculosis* strains in the 30-strain simulated dataset.

There are several factors that contribute to Strainify’s strong performance when analyzing short-read metagenomic datasets with low coverage and high genome similarity. First, its alignment-based framework provides greater tolerance for nucleotide substitutions, insertions, and deletions than k-mer–based methods, which generally depend on exact or near-exact substring matches. Consequently, k-mer approaches are more vulnerable to sequencing errors and mutations that accumulate during strain evolution, reducing their ability to detect strains with low levels of genomic divergence^31^. Second, Strainify restricts variant selection to non-recombinant regions of the core genome, which represent the most stable and evolutionarily conserved portions of a strain’s genome. These regions serve as a reliable basis for defining strain-specific variant signatures. Prior studies have shown that using variants from the accessory or recombinant genome can lead to strain misclassification, underscoring the importance of focusing on the core genome for accurate strain-level analysis^32^. Lastly, Strainify models each variant independently, an approach well-suited for short-read data, where variants in bacterial genomes are typically spaced far enough apart that most reads span at most one variant site. By incorporating all eligible SNPs and indels within the non-recombinant core genome, Strainify effectively aggregates the full set of distinguishing features between strains. This comprehensive variant representation enables Strainify to achieve maximum strain-level resolution, even in the presence of high genomic similarity. Although Strainify’s MLE framework currently focuses on biallelic variant sites, it can be readily extended to include multiallelic variant sites, which may further enhance its resolving power.

Strainify makes accessible microbiome insights at the resolution of strain-level abundance analysis. Additionally, Strainify could be coupled with taxonomic classification and metagenome assembly tools, such as Kraken 2^33^ and MetaCompass^34^, by allowing species-level abundances to be further resolved into strain-level profiles. Several key research areas stand to benefit: (1) Time-series tracking of strain-level microbiome composition, which can facilitate monitoring of disease progression and the identification of pathogenic phenotypes^35^; (2) Investigation of low-abundance microbes and their roles in shaping microbiome community structure and influencing host phenotypes^36^; and (3) Bacterial genome-wide association studies, where accurate abundance estimates are essential for linking genomic variation to phenotypic traits in microbial populations^37^.

The primary limitation of Strainify lies in its reliance on accurately assembled whole genomes as input. Low-quality genome assemblies can compromise core genome alignment, potentially leading to alignment failures or a reduction in core genome size. Furthermore, Strainify’s accuracy may be affected by increased genomic noise if the strain genomes were sequenced some time ago and have since accumulated mutations under selective pressure. Mutations at key variant sites, especially those used to distinguish closely related strains, can affect Strainify’s resolution and accuracy. This limitation can be mitigated by using strain references identified by tools such as StrainGE^38^or StrainEst^39^, or by integrating a haplotyping tool upstream when analyzing metagenomic samples with unknown strains.

As long-read sequencing gains traction due to its strengths in de novo assembly, improved mapping accuracy, and structural variant detection^40^, several tools have emerged for strain-level phasing and haplotyping. In particular, tools such as Strainy^41^ and Strainberry^42^ leverage graph-based methods to assign long reads to haplotypes. However, to our knowledge, these tools do not perform abundance estimation, and their effectiveness is often constrained by low coverage and high metagenome complexity. Although Strainify currently supports only short-read data, its high-accuracy estimation framework can potentially be extended to long reads by leveraging co-occurring variant blocks. The underlying MLE model is readily adaptable to grouped variants, offering a promising alternative to graph-based methods, particularly under conditions of low or uneven coverage and in communities with highly similar strains.

In this work, we present Strainify, a high-accuracy strain-level microbiome profiling tool that is able to both tackle the challenge presented by short-reads and low-coverage when analyzing microbial communities with bulk sequencing data. Validated on both simulated and real datasets spanning a wide range of challenging conditions, Strainify consistently outperformed StrainScan, a leading tool in the field. In a simulated 4-strain dataset with individual strain coverages as low as only 1% of the genome covered with short-read sequencing data, Strainify achieved an average absolute error three times lower than that of StrainScan. Strainify’s performance is especially notable under low sequencing coverage and high genome similarity, conditions where existing methods typically fail. By enabling robust and precise quantification of closely related strains in low-coverage settings, Strainify opens new possibilities for tracking microbial dynamics and uncovering strain-level functional variation in short-read sequenced metagenomes.

## Methods

Strainify is a pipeline of several steps that takes as input a set of assembled “reference” genomes for the particular strains of interest, and a short-read metagenomic sequencing output for a sample containing those strains and in which their relative abundance is of interest. The first steps of Parsnp are to align the reference genomes and map reads from the sample back to the alignment (by way of a representative sequence). Within the core genome (i.e., sites shared across all genomes), variant (SNP and indel) positions are identified, and for each site, the allele pattern across genomes is recorded along with read depth and allele frequency. These data collectively form the sufficient statistics for a maximum likelihood model in which the parameters are the vector of strain abundances. Strainify implements a solver for this model that outputs the solution. Individual steps of the Strainify pipeline are discussed in more detail below.

### Computing allele expression patterns and frequencies

#### Genome alignment

The first step in the pipeline is to perform a core genome alignment of the reference genomes using Parsnp^43^. This outputs a set of locally colinear blocks, which are multiple sequence alignments of particular segments of the input genomes which are identified as orthologous for all input genomes. These blocks are then scanned for sites with informative SNPs and indels, and the locations are noted. Parsnp randomly chooses one genome to be the reference, and variants are computed relative to this genome.

#### Variant calling and filtering

Variants are identified using a custom script that processes Parsnp alignment results through `wgatools`^44^ and `bcftools`^45^. Positions with exactly one variant relative to the reference base are selected. Variants are then filtered to remove recombination sites, a step that prevents the choice of reference genome from introducing biases to abundance estimates. Recombination filtering is done using a custom algorithm described under *Filtering of Variants*.

#### Read mapping and counting

Sample reads are mapped to the single reference genome using `bwa mem`^46^. The resulting alignments are processed with `samtools`^45^ to filter and index the reads, after which read depth and allele frequencies are computed at each variant locus. The resulting data, which include the allele patterns across reference genomes, observed allele frequencies, and read depths, are then integrated into the statistical model described below.

### Filtering of variants

Recombination sites, often indicated by a high density of variants, tend to confuse read mappers and lead to unmapped or mismapped reads^47–49^. Since the downstream statistical inference is highly sensitive to read mapping accuracy, a filtering step is applied to remove variants located in potential recombination regions. In the default settings, the core genome is divided into 500 base pair non-overlapping windows, and a chi-squared test is conducted to compare the variant density in each window with the average variant density of the entire core genome. The average variant density is calculated as the total number of variants in the core genome divided by the length of the core genome. Window size and window overlaps are provided as adjustable parameters. Windows with p-values below 0.05, computed using the survival function of the chi-squared distribution, are identified as potential recombination sites and excluded from downstream analysis.

The chi-squared statistic is calculated as:

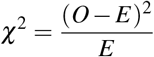

where *O* is the observed variant density in a particular window, and *E* is the average variant density of the core genome.

The p-value is computed using the survival function (right-tail probability) of the chi-squared distribution:

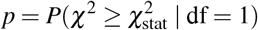

or equivalently:

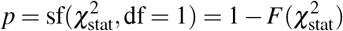

where 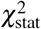 is the computed chi-squared statistic, df is the degrees of freedom, and *F* is the cumulative distribution function (CDF) of the chi-squared distribution.

### Maximum likelihood (MLE) model for relative abundance estimation

The MLE model used was adapted from a method used on samples containing multiple strains of SARS-CoV-2 together to estimate the relative strain abundances^50^.

Let *i* = 1, 2, …, *s* be the set of sites containing informative variants (SNPs and indels), *j* = 1, …, *v* be the set of genomes in the sample, and 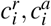 be the observed counts of the reference and alternate (allele) reads at site *i*. Let **P** = *{P*_*i j*_*}* be an (*s×v*) matrix containing the allele expression patterns, where *P*_*i j*_ is 1 if genome *j* has the allele base at variant site *i*, and 0 otherwise. Finally, let **w** = (*w*_1_, …, *w*_*v*_) be the vector of abundances for each strain, which must sum to 1.

Note in this model that for a given variant site *i*, the combined abundance of the allele strains is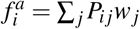, and the combined abundance of the reference strains is its complement, 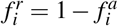 So if we let 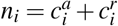 be the total number of reads mapping to that site, then the observed allele/reference count has a binomial distribution *binom*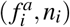 If we consider every site’s allele/reference counts as independent observations, we have the joint likelihood function:

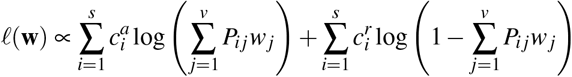

In-depth properties of this model as well as an algorithm for solving for the MLE estimate of **w** are given in the original paper on viral strains^50^. Strainify estimates strain abundances by solving the MLE problem formulated as a convex optimization task. We use CVXPY^51^, a Python-based modeling language for convex optimization to solve this problem efficiently. The default solver is ECOS^52^, which is fast and accurate for well-conditioned problems. If ECOS fails to converge, we fall back to SCS^52^, which is more robust but typically slower and less precise.

#### Entropy weighting of variants

When sequencing coverages are low, abundance estimates are more susceptible to noise caused by sequencing errors or mutations. In particular, variants which are unique to one or a small number of query genomes have a higher chance of biasing the MLE model. To increase the robustness of the model, we offer an option to weight the contribution of each variant to the likelihood function by its Shannon entropy. For a variant site *i*, the Shannon entropy (*H*_*i*_) at that site is defined as:

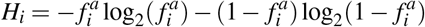

where 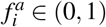is the frequency of the allele. *H*_*i*_ is maximized when half of the genomes have the 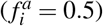allele, and it decreases as the allele frequency moves away from 0.5 in either direction. *H*_*i*_ is binary since only sites with exactly one allele are included. Thus weighting with this gives higher emphasis to sites that are more informative and less prone to variance.

The following entropy-weighted log-likelihood function is then maximized to obtain relative abundance estimates:

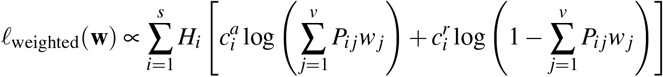

### Prediction accuracy evaluation

In experiments with simulated datasets where the ground truth abundances are known, different accuracy metrics were applied depending on the complexity of the community. For the dataset with four strains, accuracy of abundance estimates is presented as the difference from ground truth. For the dataset with 30 strains, accuracy is summarized using the Jensen-Shannon divergence^53^ between the estimated and true relative abundances. If not all queried strains were detected from a dataset, we filled in zero as the abundance for each missing strain to ensure that the true and estimated relative abundance had the same dimension. For two probability distributions *T* and *P*, their Jensen-Shannon divergence is a value between [0, 1] and is defined as^25^:

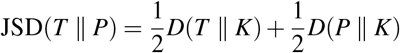

where

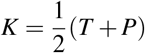

and *D*(*T*∥*K*) is called the Kullback-Leibler divergence from *T* to *K* and is defined as:

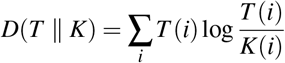

## Acknowledgements

This work was supported in part by the Texas Medical Center Genomic Center for Infectious Diseases (TMC-GCID) through NIH NIAID grants 1U19AI144297 and P01AI152999 as well as NSF awards IIS-2239114 and EF-2126387.

## Author contributions statement

Rossie S. Luo (Conceptualization, Methodology, Software, Validation, Formal analysis, Investigation, Data Curation, Writing - Original Draft, Writing - Review & Editing, Visualization, Project administration), Bryce Kille (Methodology, Software, Validation, Formal analysis, Writing - Review & Editing), Ellen E. Vaughan (Conceptualization, Writing - Review & Editing), Justin R. Clark (Conceptualization, Writing - Review & Editing), Anthony W. Maresso (Conceptualization, Writing - Review & Editing, Supervision, Funding acquisition), Michael G. Nute (Conceptualization, Methodology, Software, Validation, Formal analysis, Investigation, Writing - Original Draft, Writing - Review & Editing, Visualization, Supervision, Project administration), Todd J. Treangen (Conceptualization, Methodology, Software, Validation, Formal analysis, Investigation, Data Curation, Writing - Original Draft, Writing - Review & Editing, Visualization, Supervision, Project administration, Funding acquisition)

## Additional information

### Code availability

The source code for Strainify is available at: https://github.com/treangenlab/Strainify.

## References

1. Van Rossum, T., Ferretti, P., Maistrenko, O. M. & Bork, P. Diversity within species: interpreting strains in microbiomes. Nat. Rev. Microbiol. 18, 491–506 (2020).

2. Yan, Y., Nguyen, L. H., Franzosa, E. A. & Huttenhower, C. Strain-level epidemiology of microbial communities and the human microbiome. Genome medicine 12, 71 (2020).

3. Anderson, B. D. & Bisanz, J. E. Challenges and opportunities of strain diversity in gut microbiome research. Front. Microbiol. 14, 1117122 (2023).

4. Wassenaar, T. M. Insights from 100 years of research with probiotic e. coli. Eur. J. Microbiol. Immunol. 6, 147–161 (2016).

5. Shevchenko, S. G., Radey, M., Tchesnokova, V., Kisiela, D. & Sokurenko, E. V. Escherichia coli clonobiome: assessing the strain diversity in feces and urine by deep amplicon sequencing. Appl. Environ. Microbiol. 85, e01866–19 (2019).

6. Oles, R. E. et al. Pangenome comparison of bacteroides fragilis genomospecies unveils genetic diversity and ecological insights. Msystems 9, e00516–24 (2024).

7. Huber, A. R. et al. Improved detection of colibactin-induced mutations by genotoxic e. coli in organoids and colorectal cancer. Cancer Cell 42, 487–496 (2024).

8. Flores Ventura, E. et al. Mother-to-infant vertical transmission in early life: a systematic review and proportional meta-analysis of bifidobacterium strain transmissibility. npj Biofilms Microbiomes 11, 121 (2025).

9. Ferretti, P. et al. Mother-to-infant microbial transmission from different body sites shapes the developing infant gut microbiome. Cell host & microbe 24, 133–145 (2018).

10. Chen-Liaw, A. et al. Gut microbiota strain richness is species specific and affects engraftment. Nature 637, 422–429 (2025).

11. Ianiro, G. et al. Variability of strain engraftment and predictability of microbiome composition after fecal microbiota transplantation across different diseases. Nat. Medicine 28, 1913–1923 (2022).

12. Podlesny, D. et al. Identification of clinical and ecological determinants of strain engraftment after fecal microbiota transplantation using metagenomics. Cell Reports Medicine 3 (2022).

13. Blanco-Míguez, A. et al. Extending and improving metagenomic taxonomic profiling with uncharacterized species using metaphlan 4. Nat. biotechnology 41, 1633–1644 (2023).

14. Shen, C., Wedell, E., Pop, M. & Warnow, T. Tipp3 and tipp3-fast: Improved abundance profiling in metagenomics. PLoS computational biology 21, e1012593 (2025).

15. Ghazi, A. R., Münch, P. C., Chen, D., Jensen, J. & Huttenhower, C. Strain identification and quantitative analysis in microbial communities. J. Mol. Biol. 434, 167582 (2022).

16. Segura Munoz, R. R. et al. Experimental evaluation of ecological principles to understand and modulate the outcome of bacterial strain competition in gut microbiomes. The ISME journal 16, 1594–1604 (2022).

17. Quince, C., Walker, A. W., Simpson, J. T., Loman, N. J. & Segata, N. Shotgun metagenomics, from sampling to analysis. Nat. biotechnology 35, 833–844 (2017).

18. Debray, R., Dickson, C. C., Webb, S. E., Archie, E. A. & Tung, J. Shared environments complicate the use of strain-resolved metagenomics to infer microbiome transmission. Microbiome 13, 59 (2025).

19. Sczyrba, A. et al. Critical assessment of metagenome interpretation—a benchmark of metagenomics software. Nat. methods 14, 1063–1071 (2017).

20. Viver, T. et al. Towards estimating the number of strains that make up a natural bacterial population. Nat. Commun. 15, 544 (2024).

21. Truong, D. T., Tett, A., Pasolli, E., Huttenhower, C. & Segata, N. Microbial strain-level population structure and genetic diversity from metagenomes. Genome research 27, 626–638 (2017).

22. Luo, C. et al. Constrains identifies microbial strains in metagenomic datasets. Nat. biotechnology 33, 1045–1052 (2015).

23. Scholz, M. et al. Strain-level microbial epidemiology and population genomics from shotgun metagenomics. Nat. methods 13, 435–438 (2016).

24. Quince, C. et al. Desman: a new tool for de novo extraction of strains from metagenomes. Genome biology 18, 181 (2017).

25. Liao, H., Ji, Y. & Sun, Y. High-resolution strain-level microbiome composition analysis from short reads. Microbiome 11, 183 (2023).

26. Heber, K., Tian, S., Betancurt-Anzola, D., Koo, H. & Bisanz, J. E. StrainR2 accurately deconvolutes strain-level abundances in synthetic microbial communities. Bioinformatics 41, btaf440 (2025).

27. Huang, W., Li, L., Myers, J. R. & Marth, G. T. Art: a next-generation sequencing read simulator. Bioinformatics 28, 593–594 (2012).

28. Verma, H. et al. Genome analyses of 174 strains of mycobacterium tuberculosis provide insight into the evolution of drug resistance and reveal potential drug targets. Microb. Genomics 7, 000542 (2021).

29. Poyet, M. et al. A library of human gut bacterial isolates paired with longitudinal multiomics data enables mechanistic microbiome research. Nat. medicine 25, 1442–1452 (2019).

30. Rodola, G. Psutil documentation. Psutil. https://psutil.readthedocs.io/en/latest (2020).

31. Roberts, M. D., Davis, O., Josephs, E. B. & Williamson, R. J. k-mer-based approaches to bridging pangenomics and population genetics. Mol. biology evolution 42, msaf047 (2025).

32. Segerman, B. The genetic integrity of bacterial species: the core genome and the accessory genome, two different stories. Front. cellular infection microbiology 2, 116 (2012).

33. Wood, D. E., Lu, J. & Langmead, B. Improved metagenomic analysis with kraken 2. Genome biology 20, 257 (2019).

34. Luan, T. et al. Reference-guided assembly of metagenomes with metacompass. Cell Reports Methods (2025).

35. Fang, X. et al. Metagenomics-based, strain-level analysis of escherichia coli from a time-series of microbiome samples from a crohn’s disease patient. Front. microbiology 9, 2559 (2018).

36. Han, G., Luong, H. & Vaishnava, S. Low abundance members of the gut microbiome exhibit high immunogenicity. Gut Microbes 14, 2104086 (2022).

37. Yang, Q. et al. Bacterial genome-wide association studies: exploring the genetic variation underlying bacterial phenotypes. Appl. Environ. Microbiol. e02512–24 (2025).

38. van Dijk, L. R. et al. Strainge: a toolkit to track and characterize low-abundance strains in complex microbial communities. Genome biology 23, 74 (2022).

39. Albanese, D. & Donati, C. Strain profiling and epidemiology of bacterial species from metagenomic sequencing. Nat. communications 8, 2260 (2017).

40. Amarasinghe, S. L. et al. Opportunities and challenges in long-read sequencing data analysis. Genome biology 21, 30 (2020).

41. Kazantseva, E., Donmez, A., Frolova, M., Pop, M. & Kolmogorov, M. Strainy: phasing and assembly of strain haplotypes from long-read metagenome sequencing. Nat. Methods 21, 2034–2043 (2024).

42. Vicedomini, R., Quince, C., Darling, A. E. & Chikhi, R. Strainberry: automated strain separation in low-complexity metagenomes using long reads. Nat. Commun. 12, 4485 (2021).

43. Kille, B. et al. Parsnp 2.0: scalable core-genome alignment for massive microbial datasets. Bioinformatics 40, btae311 (2024).

44. Wei, W. et al. wgatools: an ultrafast toolkit for manipulating whole-genome alignments. Bioinformatics 41, btaf132 (2025).

45. Danecek, P. et al. Twelve years of samtools and bcftools. Gigascience 10, giab008 (2021).

46. Li, H. Aligning sequence reads, clone sequences and assembly contigs with bwa-mem. 2013 (2013).

47. Dixit, P. D., Pang, T. Y., Studier, F. W. & Maslov, S. Recombinant transfer in the basic genome of escherichia coli. Proc. Natl. Acad. Sci. 112, 9070–9075 (2015).

48. Alser, M. et al. Technology dictates algorithms: recent developments in read alignment. Genome biology 22, 249 (2021).

49. Rahman, F. et al. Benchngs: An approach to benchmark short reads alignment tools. arXiv preprint 1504.06659 (2015).

50. Valieris, R. et al. A mixture model for determining sars-cov-2 variant composition in pooled samples. Bioinformatics 38, 1809–1815 (2022).

51. Diamond, S. & Boyd, S. Cvxpy: A python-embedded modeling language for convex optimization. J. Mach. Learn. Res. 17, 1–5 (2016).

52. Gilani, A. & Liu, H. A comprehensive framework for benchmarking algorithms across hyperparameter spaces (2024).

53. Fuglede, B. & Topsoe, F. Jensen-shannon divergence and hilbert space embedding. In International symposium onInformation theory, 2004. ISIT 2004. Proceedings., 31 (IEEE, 2004).

