## Supplemental materials for "Strainify: Strain-Level Microbiome Profiling for Low-Coverage Short-Read Metagenomic Datasets"

| Strain Name | NCBI Genome Assembly |
|---|---|
| E24377A | GCF_000017745.1 |
| H10407 | GCF_000210475.1 |
| Sakai | GCF_000008865.1 |
| UTI89 | GCF_000013265.1 |

**Supplementary Table 1**. NCBI genome assemblies of *Escherichia coli* genomes used in 4-strain simulated dataset.

| NCBI Genome Assembly | Dominant | Uniform | Dirichlet |
|---|---|---|---|
| GCA_000449285.1 | 0.029669856491129316 | 0.03333333 | 0.024134860319189405 |
| GCA_004315405.1 | 0.00095145 | 0.03333333 | 0.02205481 |
| GCA_007114605.1 | 0.01963362 | 0.03333333 | 0.011622329478844272 |
| GCA_015703685.1 | 0.01291735 | 0.03333333 | 0.0500776 |
| GCA_016078175.1 | 0.00031895 | 0.03333333 | 0.00443426 |
| GCA_018736545.1 | 0.00300723 | 0.03333333 | 0.06679453 |
| GCA_018855955.1 | 0.0070101 | 0.03333333 | 0.00057861 |
| GCA_018880885.1 | 0.00145294 | 0.03333333 | 0.03925418 |
| GCA_018895495.1 | 0.00763353 | 0.03333333 | 0.04843954 |
| GCA_018908095.1 | 0.0047932 | 0.03333333 | 0.010294852820230583 |
| GCA_018942535.1 | 0.010304696190692061 | 0.03333333 | 0.09023228 |
| GCA_018962215.1 | 0.00382334 | 0.03333333 | 0.040226820890580696 |
| GCA_019763655.1 | 2.8582240988598002e-05 | 0.03333333 | 0.019818619313078926 |

| | | | |
|---|---|---|---|
| GCA_021113555.1 | 0.02169616 | 0.03333333 | 0.03834307 |
| GCA_021360505.2 | 0.00442455 | 0.03333333 | 0.046952757506868135 |
| GCA_022278845.1 | 0.00981188 | 0.03333333 | 0.00287416 |
| GCA_024260145.1 | 0.00964125 | 0.03333333 | 0.05190112 |
| GCA_028234055.1 | 0.011970214251408314 | 0.03333333 | 0.07695694 |
| GCA_029490295.1 | 0.0003669 | 0.03333333 | 0.042706428924598576 |
| GCA_030491795.1 | 0.00129693 | 0.03333333 | 0.046331763078327835 |
| GCA_032081245.1 | 0.00053881 | 0.03333333 | 0.00528316 |
| GCA_039712125.1 | 0.00350071 | 0.03333333 | 0.011075275497292193 |
| GCA_045892185.1 | 0.8 | 0.03333333 | 0.012834965740256556 |
| GCA_045892665.1 | 0.00309252 | 0.03333333 | 0.01097331 |
| GCA_046754885.1 | 0.00672132 | 0.03333333 | 0.03596072 |
| GCA_900690145.1 | 0.0024446 | 0.03333333 | 0.03630088 |
| GCA_901002505.1 | 0.00668387 | 0.03333333 | 0.02971053 |
| GCA_901003235.1 | 0.013868907003539616 | 0.03333333 | 0.01599363 |
| GCA_901006535.1 | 0.00027638 | 0.03333333 | 0.08016763 |
| GCA_901008875.1 | 0.00212016 | 0.03333333 | 0.027670372812985896 |

**Supplementary Table 2.** *Clostridioides difficile* genomes and ground truth of relative abundances used in simulated 30-strain dataset.

| **NCBI Genome Assembly** | **Dominant** | **Uniform** | **Dirichlet** |
|---|---|---|---|

| | | | |
|---|---|---|---|
| GCA_000145195.1 | 0.012627062793755748 | 0.03333333 | 0.00636025 |
| GCA_000240055.1 | 0.00533238 | 0.03333333 | 0.05523646 |
| GCA_001063605.1 | 0.00128586 | 0.03333333 | 0.00697309 |
| GCA_003384445.1 | 0.022018284217403553 | 0.03333333 | 0.03845441645673156 |
| GCA_009737065.1 | 0.00048716 | 0.03333333 | 0.0370803 |
| GCA_009737125.1 | 1.2225757654816557e-05 | 0.03333333 | 0.00033523 |
| GCA_024329005.1 | 0.00409986 | 0.03333333 | 0.09437604 |
| GCA_029338475.1 | 0.00733442 | 0.03333333 | 0.03007394 |
| GCA_040461445.1 | 0.00044713 | 0.03333333 | 0.010711457573893312 |
| GCA_040462935.1 | 0.00186593 | 0.03333333 | 0.00788163 |
| GCA_040464095.1 | 0.8 | 0.03333333 | 0.06650191 |
| GCA_040464235.1 | 0.00094263 | 0.03333333 | 0.027845306146079194 |
| GCA_040464925.1 | 0.00661177 | 0.03333333 | 0.02481018 |
| GCA_040465575.1 | 0.00350293 | 0.03333333 | 0.034440238219905786 |
| GCA_040465815.1 | 0.00013131 | 0.03333333 | 0.1229489305951106 |
| GCA_040466695.1 | 0.00259146 | 0.03333333 | 0.019118521703543848 |
| GCA_040467975.1 | 0.010786945192934307 | 0.03333333 | 0.019626421562760548 |
| GCA_040469115.1 | 0.011527724874628137 | 0.03333333 | 0.059176861056372244 |
| GCA_040470605.1 | 0.00413863 | 0.03333333 | 0.023934050963412848 |
| GCA_040470885.1 | 0.02781489 | 0.03333333 | 0.00968038 |

| | | | |
|---|---|---|---|
| GCA_040470915.1 | 0.02111688 | 0.03333333 | 0.032233667606875176 |
| GCA_040474495.1 | 0.00385304 | 0.03333333 | 0.053597461281866696 |
| GCA_040475435.1 | 0.00510018 | 0.03333333 | 0.00034806 |
| GCA_040475615.1 | 0.00251077 | 0.03333333 | 0.017201138662083337 |
| GCA_040483815.1 | 0.00333812 | 0.03333333 | 0.00720894 |
| GCA_040486575.1 | 0.00077164 | 0.03333333 | 0.050512429579582144 |
| GCA_040488775.1 | 0.015072070402719498 | 0.03333333 | 0.030681846702010258 |
| GCA_040491195.1 | 0.00215868 | 0.03333333 | 0.00133727 |
| GCA_040491815.1 | 0.01105467 | 0.03333333 | 0.09001821 |
| GCA_965137345.1 | 0.011465330327484752 | 0.03333333 | 0.021295367909334652 |

**Supplementary Table 3.** *Cutibacterium acnes* genomes and ground truth of relative abundances used in simulated 30-strain dataset.

| NCBI Genome Assembly | Dominant | Uniform | Dirichlet |
|---|---|---|---|
| GCA_001607555.1 | 0.011942240262230385 | 0.03333333 | 0.013572680735176776 |
| GCA_011955845.2 | 0.00083818 | 0.03333333 | 0.00432867 |
| GCA_012111275.1 | 0.00356569 | 0.03333333 | 0.0717291 |
| GCA_012254085.1 | 0.00749518 | 0.03333333 | 0.03997457 |
| GCA_012461225.1 | 0.00164341 | 0.03333333 | 0.1089946225808919 |
| GCA_014576175.1 | 0.00011047 | 0.03333333 | 0.010264162499644843 |
| GCA_014689615.1 | 0.014683473324232319 | 0.03333333 | 0.03764474 |

| | | | |
|---|---|---|---|
| GCA_015287005.1 | 0.00423285 | 0.03333333 | 0.00569918 |
| GCA_016851645.1 | 0.04015829 | 0.03333333 | 0.00760835 |
| GCA_017017915.1 | 0.01215574 | 0.03333333 | 0.03382751 |
| GCA_017134955.1 | 0.00587333 | 0.03333333 | 0.00385853 |
| GCA_017671065.1 | 0.00129881 | 0.03333333 | 0.04096836 |
| GCA_017673085.1 | 0.00023824 | 0.03333333 | 0.00199694 |
| GCA_019261085.1 | 3.8797684704426595e-05 | 0.03333333 | 0.04416247 |
| GCA_022217905.1 | 0.011557452577829017 | 0.03333333 | 0.016688030296726955 |
| GCA_024192735.1 | 0.0028699 | 0.03333333 | 0.00461845 |
| GCA_025034035.1 | 0.00909298 | 0.03333333 | 0.11126681091240441 |
| GCA_025259005.1 | 0.00096917 | 0.03333333 | 0.06944536 |
| GCA_025263345.1 | 0.8 | 0.03333333 | 0.00116514 |
| GCA_025619945.1 | 0.00374196 | 0.03333333 | 0.02577329 |
| GCA_029739455.1 | 0.00779337 | 0.03333333 | 0.02158793 |
| GCA_031522605.1 | 0.010370655136006377 | 0.03333333 | 0.00891438 |
| GCA_032612115.1 | 0.010401243462187662 | 0.03333333 | 0.019411681906620726 |
| GCA_032816185.1 | 0.00234337 | 0.03333333 | 0.00945889 |
| GCA_039998825.1 | 0.00951453 | 0.03333333 | 0.11588941659869094 |
| GCA_044143365.1 | 0.0021172 | 0.03333333 | 0.05536011026772 1085 |
| GCA_047595605.1 | 0.012192573165273703 | 0.03333333 | 0.05803891 |
| GCA_050648015.1 | 0.00406589 | 0.03333333 | 0.00393919 |
| GCA_946402935.1 | 0.00654492 | 0.03333333 | 0.013878359193117824 |

| | | | |
|---|---|---|---|
| GCA_965256235.1 | 0.00215007 | 0.03333333 | 0.03993417 |

**Supplementary Table 4.** *Escherichia coli* genomes and ground truth of relative abundances used in simulated 30-strain dataset.

| NCBI Genome Assembly | Dominant | Uniform | Dirichlet |
|---|---|---|---|
| GCA_000177935.2 | 0.00034813 | 0.03333333 | 0.00911662 |
| GCA_000277735.2 | 0.00244056 | 0.03333333 | 0.026210962612306463 |
| GCA_000655135.1 | 0.00881421 | 0.03333333 | 0.0003782 |
| GCA_000662665.1 | 0.012862358761624951 | 0.03333333 | 0.051566794647510114 |
| GCA_000663405.1 | 0.00123684 | 0.03333333 | 0.025055288256974433 |
| GCA_000670135.1 | 0.01120997 | 0.03333333 | 0.00065187 |
| GCA_000673815.1 | 0.011365965398819969 | 0.03333333 | 0.10670724363901596 |
| GCA_000803865.1 | 0.00280041 | 0.03333333 | 0.02625398 |
| GCA_001377835.1 | 0.00310777 | 0.03333333 | 0.0243895 |
| GCA_001383535.1 | 0.00336139 | 0.03333333 | 0.00727345 |
| GCA_001387555.1 | 0.00376298 | 0.03333333 | 0.027815978132486312 |
| GCA_001389815.1 | 0.00282627 | 0.03333333 | 0.00269188 |
| GCA_001389995.1 | 0.00187595 | 0.03333333 | 0.09541764 |
| GCA_001391755.1 | 0.00296503 | 0.03333333 | 0.00982695 |
| GCA_001852955.1 | 0.018750118964315426 | 0.03333333 | 0.023770811753617727 |
| GCA_001855915.1 | 0.016436433263741303 | 0.03333333 | 0.00685532 |
| GCA_001942815.1 | 0.001835 | 0.03333333 | 0.011959789282364916 |

| | | | |
|---|---|---|---|
| GCA_001946015.1 | 0.00301971 | 0.03333333 | 0.08795594 |
| GCA_001946875.1 | 0.01908885 | 0.03333333 | 0.012703801077856585 |
| GCA_002130845.1 | 0.00514633 | 0.03333333 | 0.00685778 |
| GCA_003393435.1 | 0.00222193 | 0.03333333 | 0.00907909 |
| GCA_003396085.1 | 0.012925244115624132 | 0.03333333 | 0.07584497 |
| GCA_004106905.1 | 0.019899366285372803 | 0.03333333 | 0.0050195 |
| GCA_004109525.1 | 0.00575625 | 0.03333333 | 0.024080735570476502 |
| GCA_004114165.1 | 0.00066025 | 0.03333333 | 0.00690658 |
| GCA_008761715.1 | 0.8 | 0.03333333 | 0.06496931 |
| GCA_014900835.1 | 0.0088614 | 0.03333333 | 0.1407938867785 9025 |
| GCA_024651365.1 | 0.0012717 | 0.03333333 | 0.03412799 |
| GCA_030285745.1 | 0.014117249943981428 | 0.03333333 | 0.02250823 |
| GCA_048076165.1 | 0.00103233 | 0.03333333 | 0.053209901368789544 |

**Supplementary Table 5.** *Mycobacterium tuberculosis* genomes and ground truth of relative abundances used in simulated 30-strain dataset.

| **NCBI Genome Assembly** | **Dominant** | **Uniform** | **Dirichlet** |
|---|---|---|---|
| GCA_000177115.1 | 0.00470645 | 0.03333333 | 0.00697889 |
| GCA_001070735.1 | 0.00126893 | 0.03333333 | 0.026200592680129864 |
| GCA_001469335.1 | 0.01284694 | 0.03333333 | 0.0357033 |
| GCA_001658525.1 | 0.014948383344897504 | 0.03333333 | 0.01817303 |

| | | | |
|---|---|---|---|
| GCA_007667225.1 | 0.01646571 | 0.03333333 | 0.00505832 |
| GCA_009897165.1 | 0.00690446 | 0.03333333 | 0.032766369630207993 |
| GCA_010140405.1 | 0.00064528 | 0.03333333 | 0.015889494368876548 |
| GCA_010145255.1 | 0.00848316 | 0.03333333 | 0.01783667 |
| GCA_010147865.1 | 0.00710378 | 0.03333333 | 0.07639258 |
| GCA_010151165.1 | 0.010560645530917089 | 0.03333333 | 0.10278353042576609 |
| GCA_010153135.1 | 0.00543617 | 0.03333333 | 0.00702638 |
| GCA_010158175.1 | 0.00923158 | 0.03333333 | 0.03459187 |
| GCA_010161595.1 | 0.00203262 | 0.03333333 | 0.024795778202286112 |
| GCA_010169415.1 | 0.00182035 | 0.03333333 | 0.09862929 |
| GCA_016808565.1 | 0.8 | 0.03333333 | 0.031007899836983682 |
| GCA_016887325.1 | 0.013454943166186423 | 0.03333333 | 0.04830308 |
| GCA_021994825.1 | 0.00410507 | 0.03333333 | 0.12181640585345083 |
| GCA_022002995.1 | 0.00891034 | 0.03333333 | 0.04648249 |
| GCA_022004555.1 | 0.00459622 | 0.03333333 | 0.014207727850804583 |
| GCA_022008135.1 | 0.00585963 | 0.03333333 | 0.018689658478553205 |
| GCA_029917425.1 | 1.4496511615694832e-05 | 0.03333333 | 0.017011486686655135 |
| GCA_029917645.1 | 0.00090913 | 0.03333333 | 0.019471475993320993 |
| GCA_030235045.1 | 0.00077755 | 0.03333333 | 0.03217162 |
| GCA_030716625.1 | 0.00648055 | 0.03333333 | 0.0713048 |

| | | | |
|---|---|---|---|
| GCA_031734655.1 | 0.00941543 | 0.03333333 | 0.024283889432295824 |
| GCA_032471895.1_ASM3247189v1_genomic.fna | 0.0024157 | 0.03333333 | 0.024694281331648395 |
| GCA_037698885.1_ASM3769888v1_genomic.fna | 0.00862098 | 0.03333333 | 0.013803182210426552 |
| GCA_045501355.1_ASM4550135v1_genomic.fna | 0.00338131 | 0.03333333 | 0.00222929 |
| GCA_050578175.1_ASM5057817v1_genomic.fna | 0.00298089 | 0.03333333 | 0.00446826 |
| GCA_964209235.1_FFDD_204.1_genomic.fna | 0.02562328 | 0.03333333 | 0.00722835 |

**Supplementary Table 6.** *Staphylococcus epidermidis* genomes and ground truth of relative abundances used in simulated 30-strain dataset.

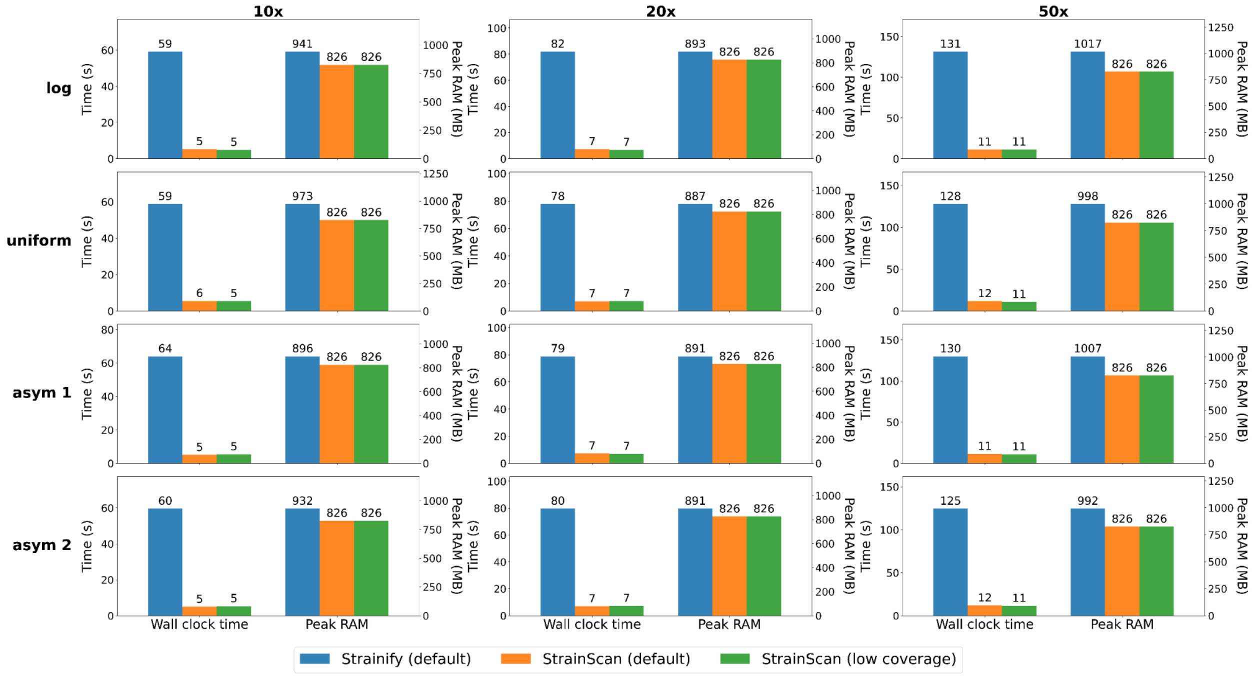


**Supplementary Figure 1.** Wall clock time (s) and peak RAM (MB) of *compute abundances* step in Strainify (default), StrainScan (default) and StrainScan (low coverage) in the 4-strain *E. coli* simulated dataset. Abundance estimates for the same dataset are presented in Figure 2 of the main text.

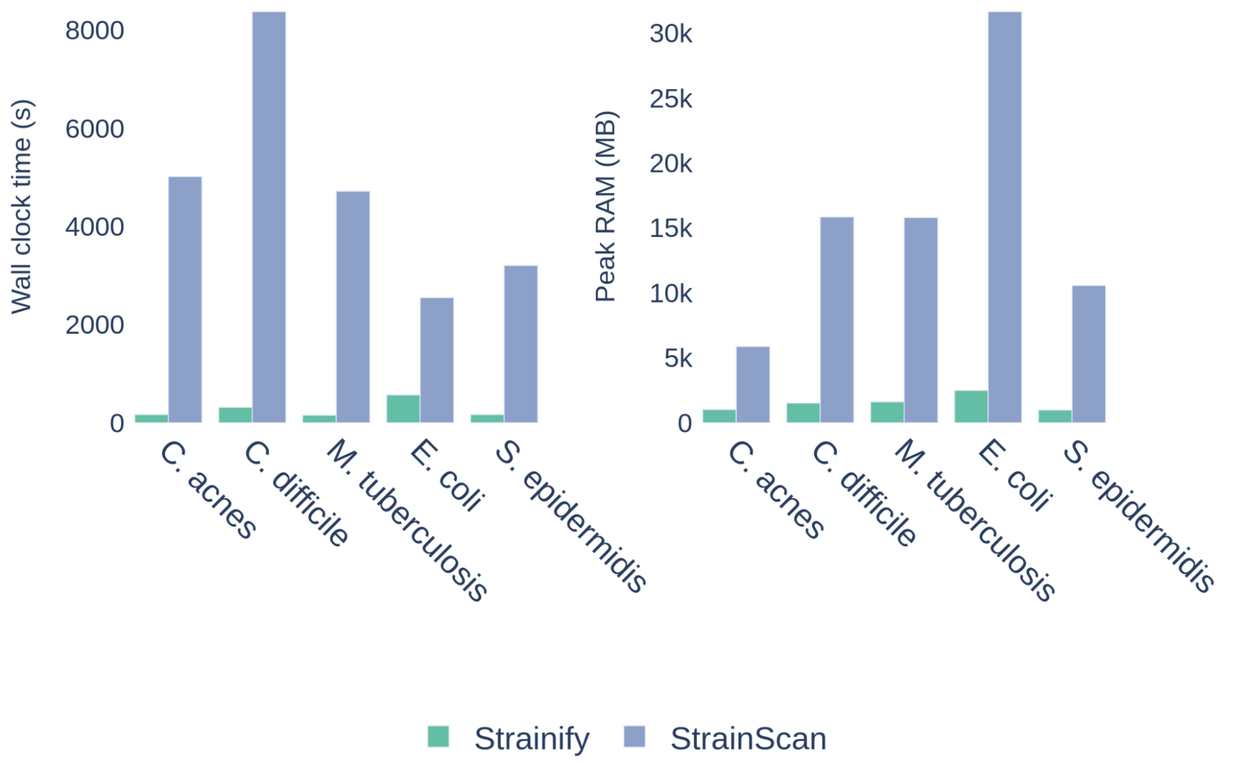


**Supplementary Figure 2.** Wall clock time (s) and peak RAM (MB) of *build variant matrix* step in Strainify and *build database* step in StrainScan in the 30-strain simulated dataset. Abundance estimates for the same dataset are presented in Figure 3 of the main text.

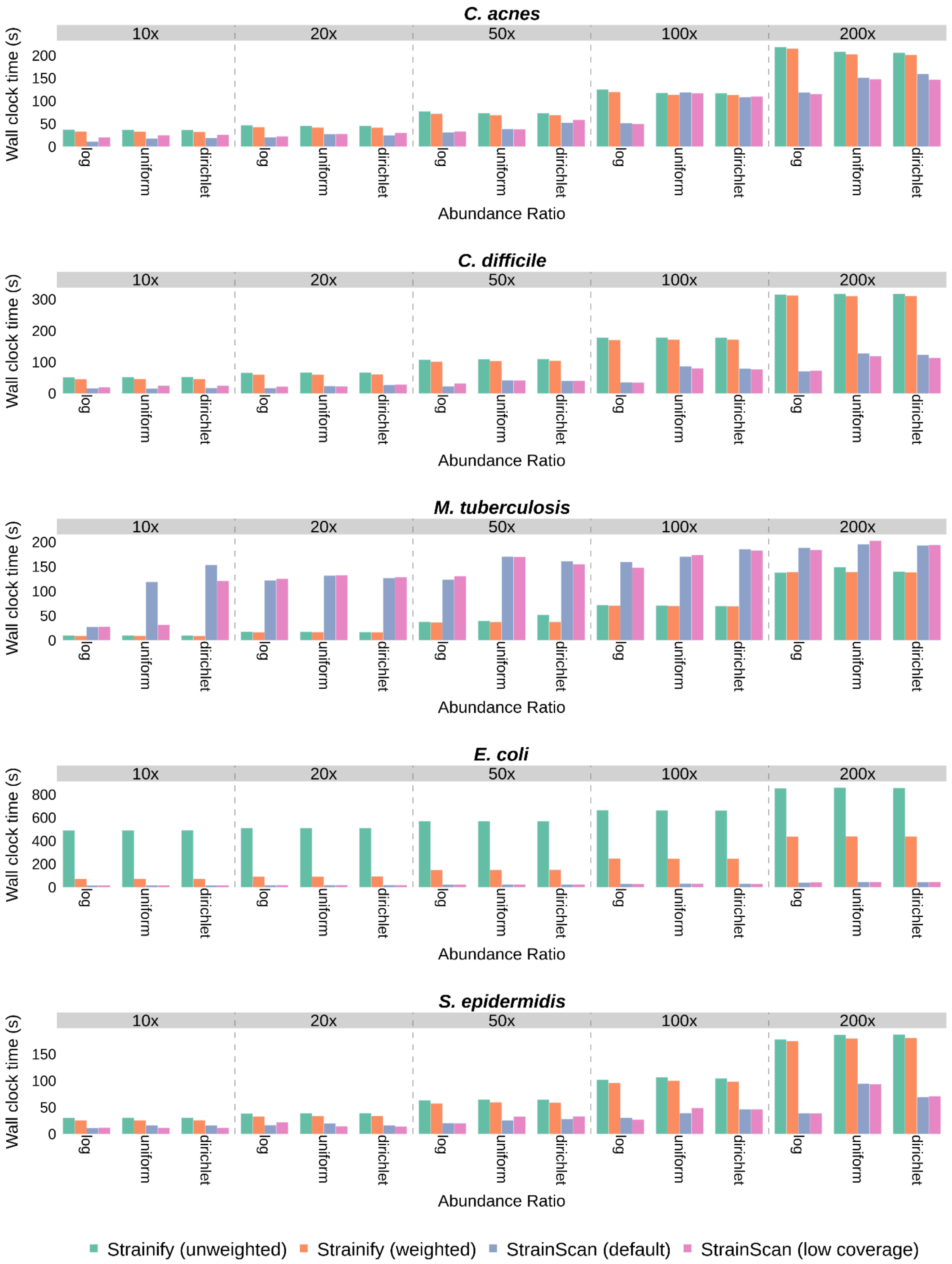


**Supplementary Figure 3.** Wall clock time (s) of the *compute abundances* step in Strainify (unweighted), Strainify (weighted), StrainScan (default) and StrainScan (low coverage) in the

30-strain simulated dataset. Abundance estimates for the same dataset are presented in Figure 3 of the main text.

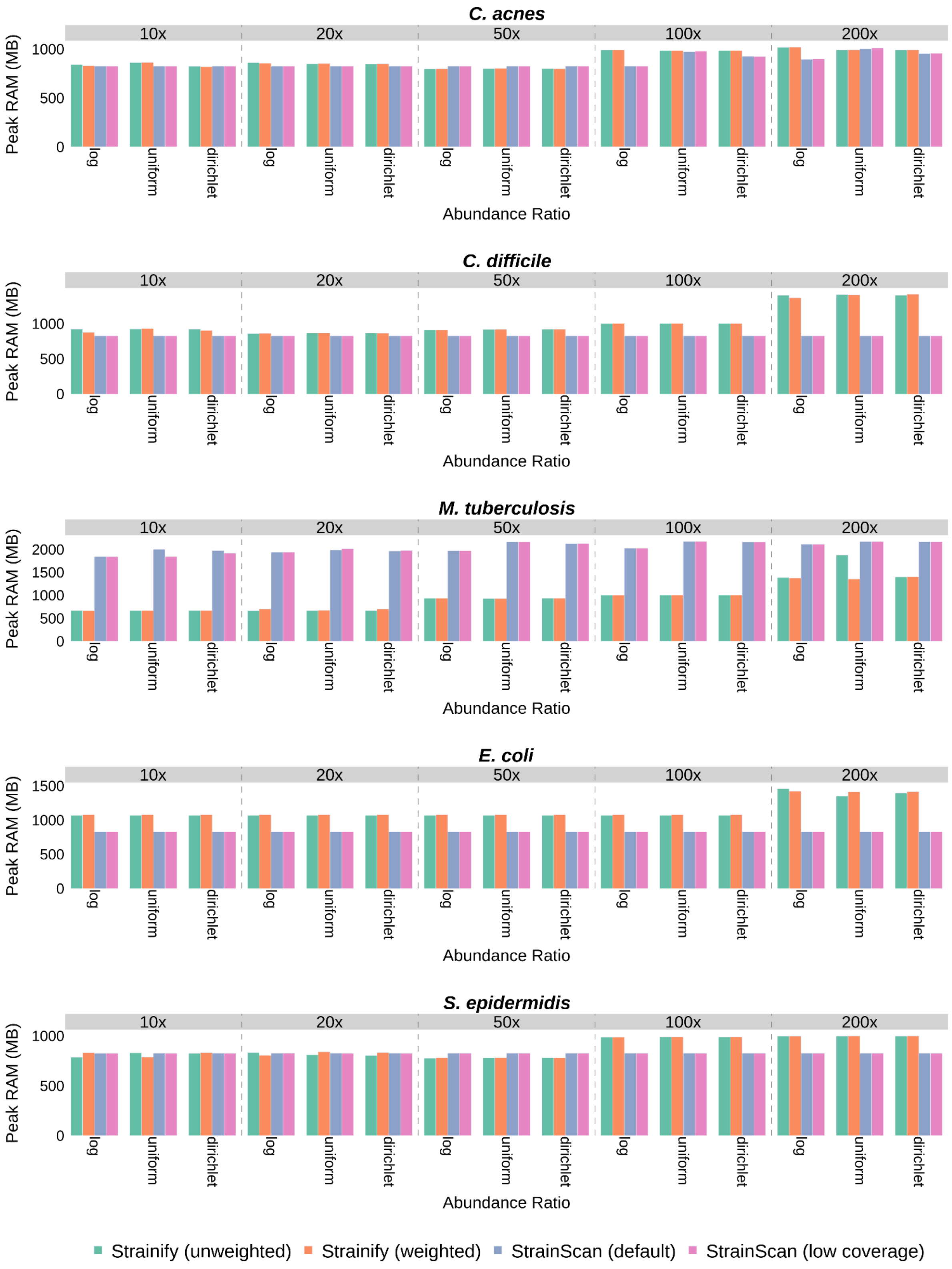

**Supplementary Figure 4.** Peak RAM (MB) of the *compute abundances* step in Strainify (unweighted), Strainify (weighted), StrainScan (default) and StrainScan (low coverage) in the 30-strain simulated dataset. Abundance estimates for the same dataset are presented in Figure 3 of the main text.
